# A truncated HIV Tat demonstrates potent and specific latency reversal activity

**DOI:** 10.1101/2023.03.02.530914

**Authors:** Ellen Van Gulck, Marion Pardons, Erik Nijs, Nick Verheyen, Koen Dockx, Christel Van den Eynde, Emilie Battivelli, Jerel Vega, Eric Florence, Brigitte Autran, Nancie M. Archin, David M. Margolis, Kristine Katlama, Chiraz Hamimi, Ilse Van den Wyngaert, Filmon Eyassu, Linos Vandekerckhove, Daniel Boden

## Abstract

A major barrier to HIV-1 cure is caused by the pool of latently infected CD4 T-cells that persist under combination antiretroviral therapy (cART). This latent reservoir is capable of producing replication-competent infectious virus once prolonged suppressive cART is withdrawn. Inducing the reactivation of HIV-1 gene expression in T-cells harboring a latent provirus in people living with HIV-1 under cART will likely result in depletion of this latent reservoir due to cytopathic effects or immune clearance. Studies have investigated molecules that reactivate HIV-1 gene expression but to date no latency reversal agent has been identified to eliminate latently infected cells harboring replication-competent HIV in cART treated individuals. Stochastic fluctuations in HIV-1 *tat* gene expression have been described and hypothesized to allow the progression into proviral latency. We hypothesized that exposing latently infected CD4+ T-cells to Tat would result in effective latency reversal. Our results indicate the capacity of a truncated Tat protein and mRNA to reactivate HIV-1 in latently infected T-cells *ex vivo* to a similar degree as the protein kinase C agonist: Phorbol 12-Myristate 13-Acetate, without T-cell activation nor any significant transcriptome perturbation.

## Introduction

The introduction of combination antiretroviral therapy (cART;(Smiley et al., 2021)) greatly increased life expectancy and improved the quality of life of people living with HIV, transforming HIV infection from a lethal to a chronic infection under cART (Deeks et al., 2013). However, cART does not cure HIV infection due to the existence and persistence of a latent viral reservoir (Chun et al., 1998, Finzi et al., 1997, Wong et al., 1997) that is established early after infection(Colby et al., 2018, Henrich et al., 2017, Whitney et al., 2014). The latent HIV reservoir is mainly formed as a small pool of long-lived memory CD4^+^ T-cells harboring an integrated latent provirus(Finzi et al., 1999, Wong et al., 1997). As the mean half-life of this latent reservoir is approximately 44 months in cART treated individuals, complete viral elimination would require 70 years of continuous cART (Finzi et al., 1999, Siliciano et al., 2003, Crooks et al., 2015). HIV cure would result in significant health benefits by removing the substantial burden of daily treatment in addition to lifting the social stigma associated with HIV.

Several experimental approaches in HIV-infected individuals have been described to decrease the size of the HIV latent reservoir to potentially allow the discontinuation of cART without the risk of viral rebound: early cART administration (Ananworanich et al., 2015), myeloablative therapy for malignancy followed by transplantation of cells lacking co-receptors for virus (Hutter et al., 2009), vaccine immunotherapy (Fidler et al., 2020, Leth et al., 2016), block and lock strategy, which aims to permanently silence integrated provirus by using HIV latency-promoting agents (Castro-Gonzalez et al., 2018), and the use of latency reversal agents (LRAs)in the presence of cART to induce latency reversal and clearance of the HIV virus (Archin et al., 2012, Elliott et al., 2014, Rasmussen et al., 2014).

Various classes of LRAs have been tested ex vivo individually or in combination in either immortalized CD4+ T-cell lines or primary CD4 cells from HIV-infected individuals. Examples of candidate LRAs include protein kinase C (PKC) agonists which activate multiple signaling pathways including the nuclear factor kB (NF-kB) pathway (Beans et al., 2013, Bui et al., 2017, Bullen et al., 2014, Jiang and Dandekar, 2015, Marsden et al., 2017, Pandelo Jose et al., 2014, Perez et al., 2010, Walker-Sperling et al., 2015, Williams et al., 2004), histone deacetylase (HDAC) inhibitors which induce HIV gene expression by facilitating a more transcription permissive chromatin environment, bromodomain inhibitors (Archin et al., 2012, Barton et al., 2014, Gunst et al., 2019, Laird et al., 2015, Rasmussen et al., 2014, Sogaard et al., 2015), TLR agonists (Lim et al., 2018, Offersen et al., 2016, Tsai et al., 2017, Vibholm et al., 2019), AKT agonists (Elliott et al., 2015, Kula et al., 2019, Spivak et al., 2014, Xing et al., 2011), SMAC mimetics (Nixon et al., 2020) and immunomodulators (Darcis et al., 2015, Laird et al., 2015, Spivak and Planelles, 2018, Van der Sluis et al., 2020). In initial clinical trials these LRAs alone or in combination, enhanced cell-associated HIV RNA levels, but none of these compounds led to a significant decrease in the size of the HIV latent reservoir in people living with HIV (reviewed in(Rodari et al., 2021, Kula-Pacurar et al., 2021)). As follow-up shock and kill strategies, PKC agonists or immunomodulators were combined with other interventions such as therapeutic vaccination or broadly neutralizing antibodies (bNAb). These therapies have shown promising results in non-human primates leading to the reduction of the viral reservoir or extended periods of aviremia (Borducchi et al., 2016, Borducchi et al., 2018, Halper-Stromberg et al., 2014). Recently, Kim et al showed that a single administration of the PKC modulator SUW133 combined with allogenic human peripheral blood NK cells delayed viral rebound after treatment interruption in humanized mice infected with HIV (Kim et al., 2022). There remains a need to enhance our understanding of the biology of the latent reservoir as well as to develop novel LRAs that are potent, specific, and tolerable for repeated administration.

HIV transcription is driven by its native 5’-long terminal repeat (LTR) promoter whereby transcriptional activity is auto-induced by the HIV-1 transactivator of transcription (Tat) protein leading to a powerful positive feedback loop (Karn, 1999). The transactivation capacity of Tat was reported shortly after the discovery of HIV showing a 200- to 300-fold LTR stimulation of integrated proviral transcription (Sodroski et al., 1985a, Sodroski et al., 1985b, Sodroski et al., 1985c). HIV Tat recruits the host positive transcription elongation factor b (P-TEFb) along with multiple additional transcription factors to a stem-loop RNA structure called transactivation response region (TAR) (Zhou et al., 1998) which is located immediately downstream of the LTR transcription start site. The P-TEFb complex is formed of the CyclinT1 and CDK9 subunits whereby the latter promotes the phosphorylation of the RNA Polymerase II (RNAPII) C-terminal domain and additional regulatory proteins (Fujinaga et al., 2004, Ivanov et al., 2000, Kim et al., 2002), leading to the full activation of polymerase processivity and elongation of HIV transcripts.

Several studies have shown that different levels of Tat expression can define the fate of HIV-infected cells by either producing infectious particles or establishing latency (Jordan et al., 2001). It was shown that the introduction of exogenous Tat was able to induce latency reversal both in a latently infected cell line model (Donahue et al., 2012) as well as in memory CD4+ T-cells from cART-treated individuals *in vitro* (Jordan et al., 2001, Lin et al., 2003), independently of the chromatin environment surrounding the integration site (Jordan et al., 2001). While intracellular expression of Tat prevents latency establishment, HIV infection with an attenuated Tat virus increases the frequency of latently infected cells (Pearson et al., 2008). Stochastic fluctuations of Tat expression might act as a molecular switch between active transcription and latency (Burnett et al., 2009, Weinberger et al., 2005, Weinberger et al., 2008).

The HIV-derived accessory protein Tat (86-101 aa) is translated from two different exons where exon 1 (1-72 aa) contains all the domains essential for trans-activation (Jeang, 1998). Here we seek to reduce the size of Tat to the minimum protein domains that are critical for its reactivation capacity by deleting regions that upon non-specific cell receptor binding may activate off-target signaling cascades. We explored various deletion mutants of Tat as potential candidates for latency reversal, both in cell lines and in primary CD4+ T-cells from cART-treated HIV-1 infected individuals. If low level of Tat expression plays a role in the establishment of HIV latency, we hypothesized that providing sufficient Tat protein in trans would result in efficient HIV-1 reactivation. In this study, we identified a Tat deletion mutant (T66) to be as efficient as full-length Tat protein to induce latency reversal *in vitro* and *ex vivo*. Importantly, the mutant did not induce global T-cell activation in primary CD4+ T-cells, nor did it lead to any significant perturbation of the human T-cell transcriptome. Furthermore, we investigated delivery of T66 mRNA to CD4+ T-cells via lipid nanoparticles and observed significantly enhanced HIV reactivation when compared to the T66 protein. The enhanced activity of T66 mRNA delivered by nanoparticles is likely due to more efficient cytosolic delivery as opposed to T66 protein transduction which depends on endocytic uptake mechanisms and endosomal escape of the entrapped protein.

## Results

### Identifying minimal HIV Tat activation domain

A deletion exercise was performed to delineate the essential Tat amino acid sequences that would result in the same HIV-LTR activation capacity as the 72 aa Tat exon 1 protein. The intention was that reducing the overall size of HIV Tat will: (i) remove potential anti-HIV Tat antibody epitopes, (ii) improve cellular uptake, (iii) diminish off-target effects due to reduced non-specific binding, and (iv) facilitate chemical polypeptide synthesis. The latter was chosen over bacterial or yeast recombinant protein production to exclude any TLR ligand contamination in the Tat protein. Several Tat deletion mutants, ranging from 57 aa up to 86 aa, were introduced into pDNA expression vectors. These constructs were used to co-transfect HEK293 cells along with pLTR-FLuc and pEF-RLuc reporter plasmids. Tat-86 and Tat-72 showed similar transactivation activity confirming the reported finding that exon-1 [72 aa] contains all elements essential for transactivation(Arya et al., 1985, Muesing et al., 1987). The Tat variant comprising only 66 amino acids was identified to have comparable latency reversal activity as the full-length exon Tat-72 construct whereas all smaller variants showed reduced reactivation activity (Fig. 1). This work further explored the latency reversal potential of the T66 variant in different *in vitro* and *ex vivo* latency reactivation assays.

**Figure 1:**
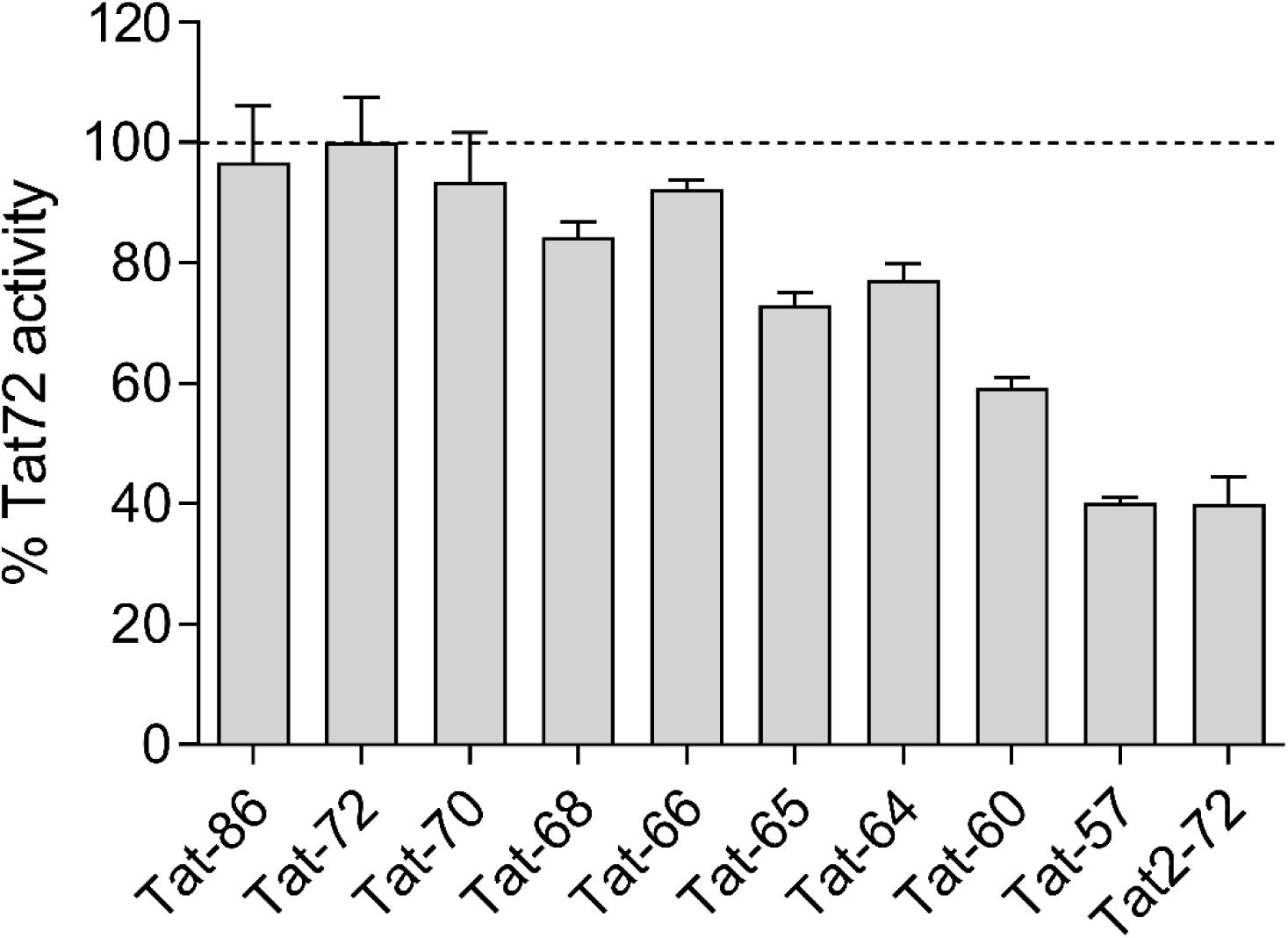
Identifying minimal Tat construct that retains maximal transactivation capacity. HEK293 T-cells were co-transfected with different Tat expression plasmids (on x-axis), pLTR-Fluc and pEF-Rluc plasmids. LTR activity was assessed via the firefly luciferase read-out along with signal normalization by Renilla luciferase. Data are normalized to LTR activity of Tat72 construct (exon1) which contains all the domains essential for transactivation. Replicate number n ≥4. Mean, error bars indicate SEM.

### Strong and consistent T66 protein mediated LTR activation of latent HIV in cell lines and primary CD4 T-cells

We first evaluated latency reversal following a dose-response with the T66 protein (ranging from 5.1 to 9 µM) in three different cell lines: MT-4-LTR-GFP, Jurkat clone 50 and Jurkat clone 32 (Fig. 2A). The percentage of GFP positive cells reached a plateau at a concentration of 6.2 µM in the three cell lines, with 87%, 63% and 51% of GFP positive cells in the MT-4-LTR-GFP, Jurkat clone 50 and Jurkat clone 32, respectively. To confirm these results in a more physiologically relevant model, a transient HIV reactivation assay was established in primary CD4+ T-cells. This assay is based on the co-infection of isolated CD4+ T-cells from HIV-seronegative donors with an HIV reporter construct and a virus expressing two antiapoptotic proteins (MCL1 and Bcl2). Following the establishment of quiescent proviral DNA and the subsequent overnight incubation of these CD4+ T-cells with PBS, T66 protein or PMA, substantial levels of reactivation were obtained with T66 and PMA in all donors tested, expressed as % increase in GFP compared to PBS-treated control (Fig. 2B). Notably, the LTR reactivation of T66-treated cells exceeded the LTR reactivation achieved by PMA stimulation in 5 out of 10 donor samples.

**Figure 2:**
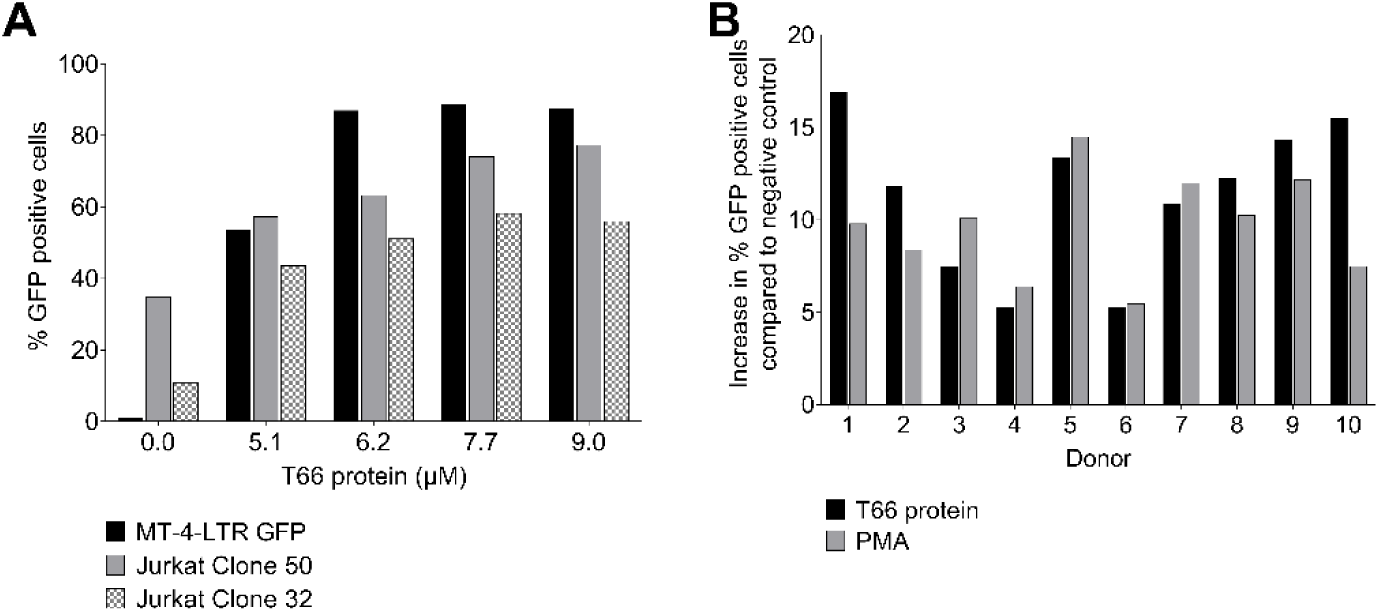
Evaluation of the capacity of the T66 protein to activate LTR in different engineered latently infected T-cell lines and CD4+ reporter cells. (A) Three different HIV-LTR GFP reporter T-cell lines: MT-4-LTR GFP (black bars), Jurkat Cl50 (grey bars) and Jurkat Cl32 (patterned bars) were incubated for 16 h with different concentrations of T66 protein (x-axis). LTR activation in these cell lines resulted in GFP expression, which was measured by flowcytometry. Percentage of GFP positive cells is plotted on the y-axis. (B) Primary CD4+ T-cells from 10 different HIV-seronegative donors (x-axis) were isolated and co-infected with an HIV dual reporter virus and a lentivirus expressing pro-survival proteins. Following infection, latently infected cells were sorted and exposed for 24h to PMA (light bars) or T66 protein (dark bars). The increase in percentage GFP positive cells compared to negative (PBS-treated cells) is plotted on the graph.

### Strong and consistent latency reversal activity in CD4 T-cells from cART-treated individuals

Next, the latency reversal capacity of the T66 protein was investigated on resting CD4+ T-cells from HIV-infected individuals under cART in four independent studies applying various methods to measure HIV latency reversal. The first two studies (Fig. 3A/B) measured cell-associated HIV RNA by RT-qPCR. HIV intracellular RNA was evaluated 24h after the addition of the compound. T66 treatment resulted in a significant increase of cell-associated RNA compared to non-stimulated controls (n=10; p=0.0002 negative control versus T66) surpassing HIV RNA levels achieved by PMA stimulation in all ten participants (Fig. 3A) (n=10; medians: 242, 2180, 422 gag RNA copies in the non-stimulated, T66 and PMA-treated cells, respectively). In the second study, T66 treatment increased the levels of cell-associated RNA compared to the non-stimulated condition in CD4 T-cells from 3 out of 4 donors. The cell-associated RNA levels were comparable to the levels observed following PHA stimulation (Fig. 3B). To evaluate virion release due to HIV-1 reactivation, p24 secretion in the supernatant was assessed by the ultrasensitive SIMOA (Fig. 3C and E). CD4+ T-cells were isolated from five cART-treated individuals and were incubated for 10 days with either T66 protein or anti-CD3/-CD28 antibodies. Both treatments led to similar p24 induction in all five donors. An additional study using a similar methodology confirmed T66-mediated p24 induction in CD8 depleted PBMC collected from five HIV-infected individuals following a 12-day stimulation (Fig. 3E). Extracellular p24 was induced in all samples after the addition of T66 protein resulting in similar or increased p24 levels with respect to CD3/CD28 treated samples. Due to donor variability peak production differs as well as time to reach maximum virus release. Finally, the quantitative viral outgrowth assay (qVOA) was applied to investigate the capacity of the T66 protein at inducing the expression of replication competent proviruses (n=6 cART-treated individuals, Fig. 3D). In this assay T66 treatment showed significant donor-to-donor variability in its ability to induce outgrowth of replication-competent virus. To conclude, T66 protein is a promising candidate for the reversal of latency in CD4+ T-cells of cART-treated individuals.

**Figure 3:**
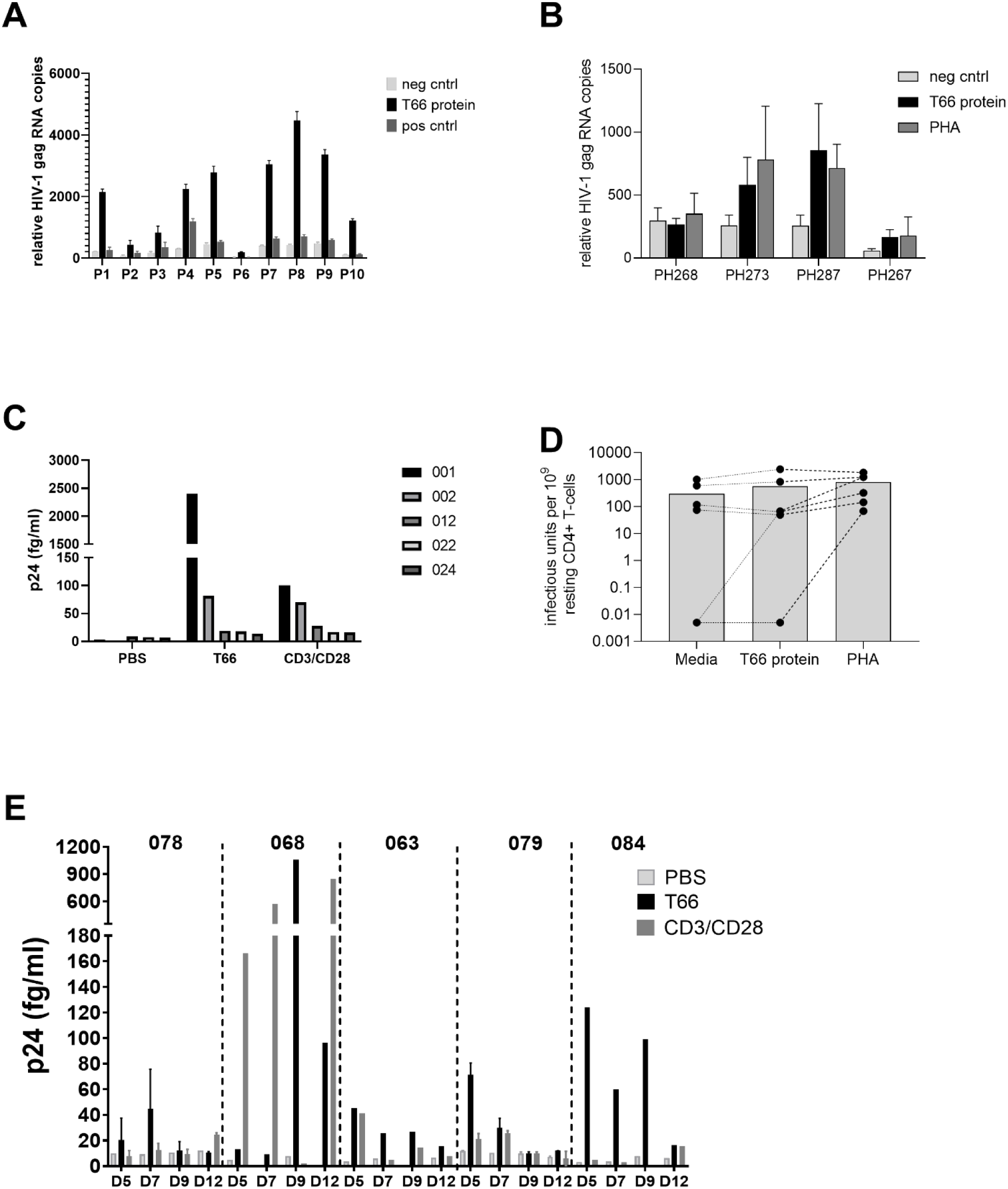
T66 protein reactivates latently infected CD4+ T-cells as measured by cell-associated RNA, p24-SIMOA, and QVOA. (A)CD4+ T-cells from 10 cART treated donors were incubated for 24h with T66 protein (black bars), PBS (light grey bars), or PMA (grey bars). HIV-1 reactivation following latency reversal was assessed by RT-qPCR. Relative HIV-1 gag RNA copies are plotted on the y-axis (15-40 replicates mean +/-SD). (B) Resting CD4+ T-cells from different HIV-1 cART treated donors were incubated for 24h with PBS (light grey bars), T66 protein (black bars) or PHA (grey bars). HIV-1 reactivation following latency reversal was assessed by RT-qPCR. Relative HIV-1 gag RNA copies are plotted on the y-axis (20replicates mean +/−SD). (C) CD4+ T-cells from 5 cART treated individuals (bars) were incubated with T66 protein, CD3CD28, or PBS. Supernatant was collected on day 10 and p24 production was measured by SIMOA technology. (D)Resting CD4+ T-cells from 5 cART treated donors were plated in replicate (n=8) at limiting dilutions of 2.5, 0.5, 0.1 and 0.025 million cells per well, activated with PHA, T66 protein or PBS in presence of allogeneic irradiated PBMCs. Culture supernatants were harvested on day 15 and assayed for virus production by p24 antigen capture ELISA. A maximum likelihood method was used to estimate the frequency of replication-competent proviruses, reported as infectious units per million of CD4+ T-cells (y-axis). (E) CD8 depleted PBMC from 5 cART treated individuals were stimulated with PBS (light grey bars), T66 protein (black bars) or CD3/CD28 (dark grey bars). Supernatant was collected on day 5, 7, 9 and 12. p24 production was measured by SIMOA technology.

### T66 does not induce global T cell activation or modifications in the CD4+ T-cells transcriptome

Clinically acceptable LRAs should not induce global T-cell activation(Prins et al., 1999). In contrast to the well-known mitogen PHA, T66 protein did not induce global T cell activation, as shown by the absence of CD69 (early activation marker) and CD25 (late activation marker) up-regulation (Fig. 4A). Moreover, micro-array experiments were conducted on total RNA extracted from CD4+ T-cells from four cART-treated individuals to study the impact of the T66 protein on the transcriptome of the cells. PBS and PHA-treated samples were included as negative and positive controls, respectively. To ensure that T66 was present and active in the HIV-infected CD4+ T-cells, latency reversal was confirmed in those samples by HIV RNA qPCR (supplementary Fig. 1). The transcriptome profile from microarray experiments revealed that T66-treated cells cluster with PBS-treated cells for all participants, suggesting no significant impact of T66 treatment on the host transcriptome (Fig. 4B). This is in sharp contrast with PHA-stimulated cells which formed a second cluster, confirming the profound impact of the mitogen PHA on the human transcriptome. Based on Volcano plots, we identified 8 genes as differentially expressed between PBS-treated samples and T66-treated samples (adjusted p<0.05, Fig. 4C). However, for these 8 differentially expressed genes, the log2 fold change was lower than 2, indicating that treatment with the T66 protein minimally perturbs the transcriptome of CD4+ T-cells.

**Figure 4:**
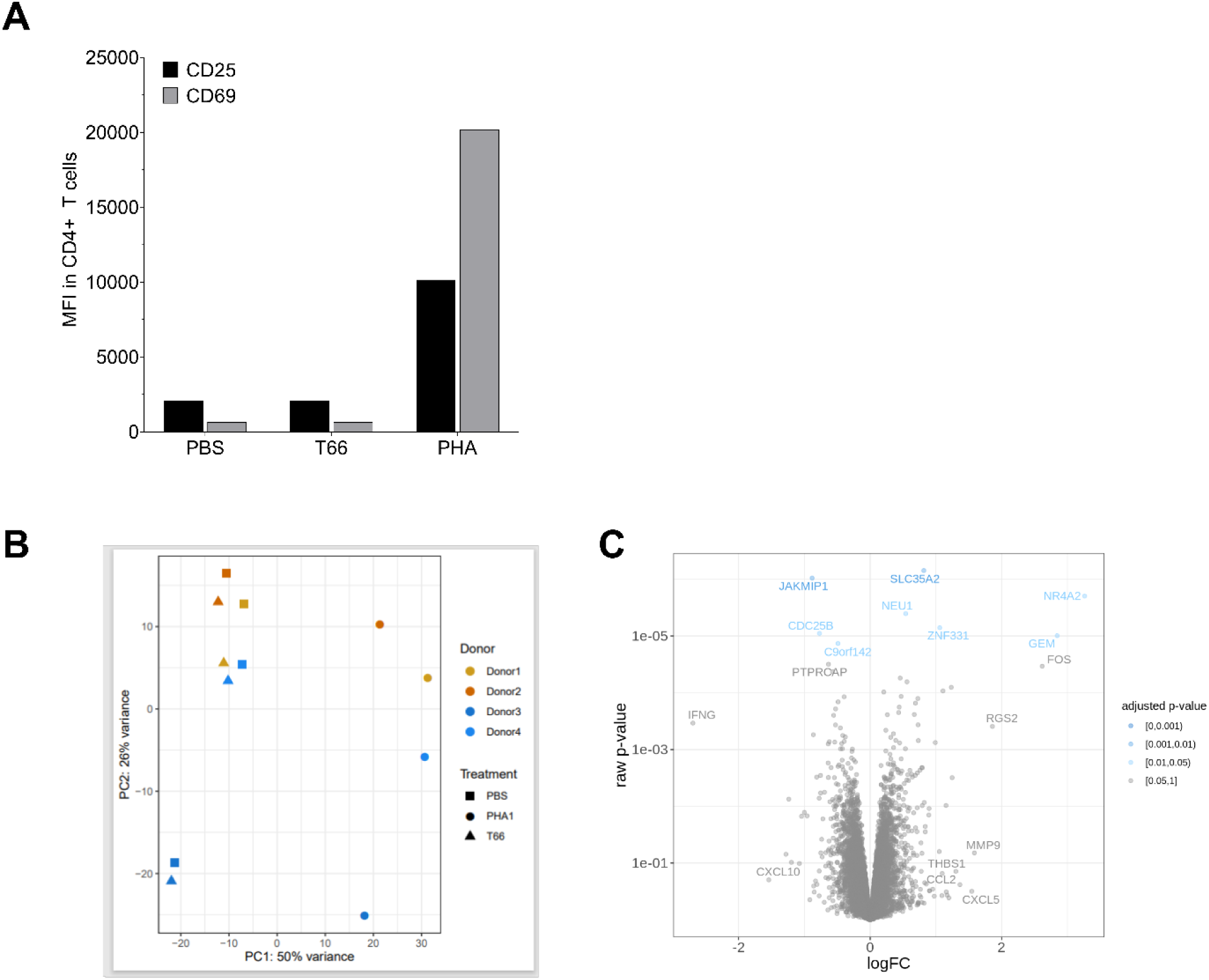
T66 protein does not induce activation of T-cells and has no effect on transcriptome. (A)CD4+ T-cells from 4 cART treated individuals were exposed overnight to PBS, T66 protein, or PMA. Expression of CD25 and CD69 was assessed by flowcytometry. The mean fluorescence intensity (MFI) is plotted for CD69 (grey bars) and CD25 (black bars). (B) 6h after stimulation cells were lysed to isolate RNA and to evaluate effects on transcriptome by microarray. Principal component analysis (PCA) plot is shown for 4 donors and 3 different treatments (PBS, T66, PHA). (C) Volcano plot comparing differentially expressed genes between PBS treated and T66 treated CD4+ T-cells.

### T66 mRNA exceeds the latency reversal levels achieved with T66 protein in T-cell lines

T66 protein showed robust latency reversal in multiple assays performed with different methodologies and in different laboratories, while neither global T-cell activation nor any significant transcriptome perturbation was observed. However, significant amounts of protein will be required for in vivo administration of T66 protein to achieve maximum HIV activation, due to limitations in cellular protein transduction and substantial endosomal trapping of the T66 protein. One strategy to address this potential liability is to use T66 mRNA delivered by lipid nanoparticles which would result in enhanced cytosolic protein exposure compared to externally applied T66 protein. Initially, a variety of commercially available reagents were evaluated for the transfection of T66 mRNA into the T-cell lymphoma MT-4-LTR-GFP cell line. Among all the different methods tested jetMessenger was the most efficient transfection reagent (data not shown). Next, the latency reversal capacity of jetMessenger delivered T66 mRNA was compared with the administration of the T66 protein (Fig. 5A). The performed dose response experiment showed that T66 mRNA induced LTR activation as measured by GFP expression reached an upper plateau at the lowest concentration tested (0.5 nM), whereas T66 protein treatment required 4 µM to achieve maximum activation. We also evaluated various lipid nanoparticle (LNP) formulations for efficient delivery of the T66 mRNA and identified LNP-2 as a promising LNP formulation (data not shown). The LNP-2 formulation was compared to jetMessenger formulation in the Jurkat clone 50 cell line (Fig. 5B). While 6.1nM jetMessenger/T66 mRNA led to a similar GFP induction as PMA (67% vs 69 % GFP(+)), 1.8nM LNP2-T66 mRNA enhanced the population of GFP (+) cells to 91.5 %. Given that the clone 50 cell line was generated via multiple rounds of PMA-stimulated GFP (+) cell sorting it was striking to see that the LNP-T66 mRNA delivery exceeded PMA-induced GFP activation. It is equally remarkable that the lowest tested LNP-2-T66 mRNA concentration (7pM) resulted in similar activation (68.4%) as the jetMessenger -T66 mRNA formulation (67.1%) at its highest concentration (6.1nM) (Fig. 5B). This translates to a potency increase of the LNP-2 formulation over the jetMessenger reagent by more than 800-fold.

**Figure 5:**
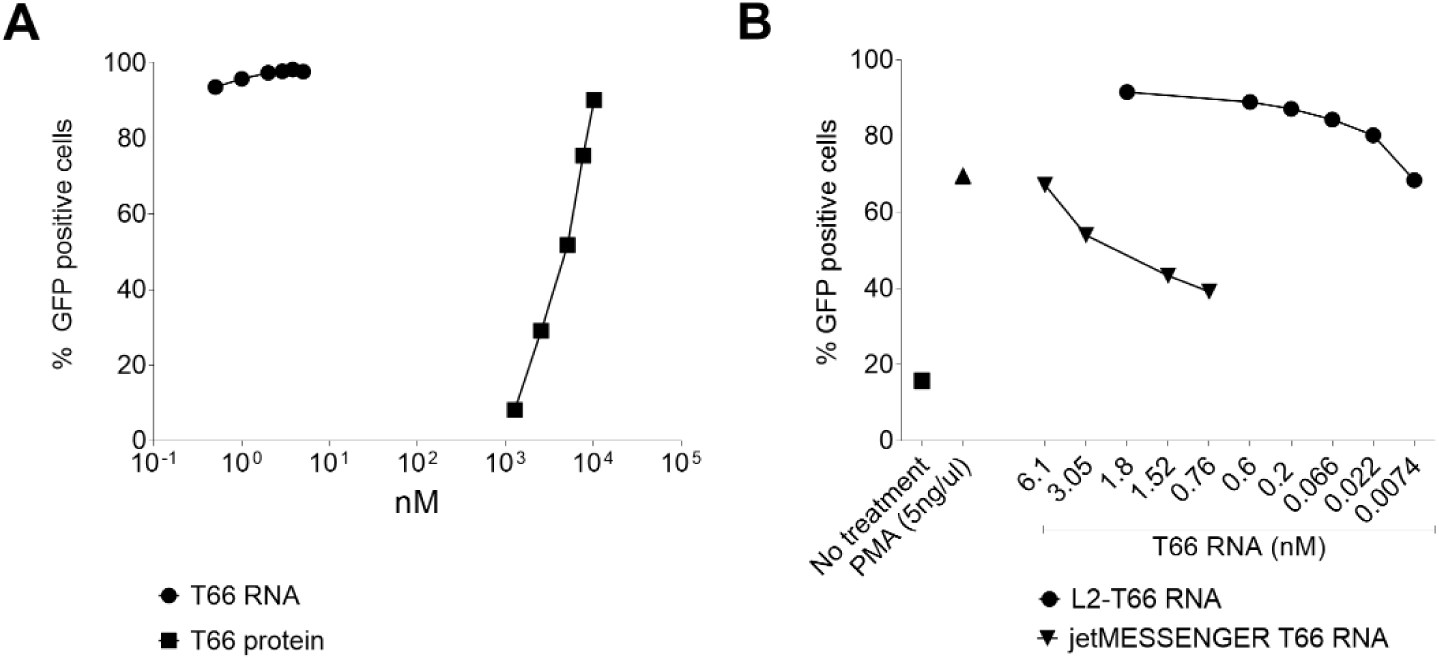
Reactivation of latently infected cell lines by T66 RNA. (A) MT-4LTR-GFP cell lines were treated with a dose response of T66 mRNA (circles) delivered by jet MESSENGER or with a dose response of T66 protein (squares). 24h post stimulation LTR activation was assessed by measuring the percentage of GFP positive cells. Fold change compared to untreated MT-4 LTR-GFP cells is plotted. (B) In Jurkat cl50 cells, two delivery methods of T66RNA were evaluated: jet MESSENGER (triangle) or LNP-2 (circles) to reactivate. LTR activation is expressed as percentage of GFP positive cells.

### LNP delivery of the T66 mRNA induces potent HIV reactivation in CD4+ T-cells from cART-treated individuals

To confirm our findings obtained in cell lines in a more physiologically relevant model, we tested the latency reversal capacity of the T66 mRNA when delivered through the LNP-2 formulation (LNP-2-T66 RNA) in CD4+ T-cells from eight cART-treated individuals. A 48h-stimulation with PMA (10nM) was used as positive control, while two negative controls were used: (i) non-stimulated cells, and (ii) a lipid nanoparticle containing the hemagglutinin A (HA) peptide from influenza. Reactivation capacity was assessed by (i) droplet digital PCR to quantify the levels of multiply spliced tat/rev mRNAs (Fig. 6A) and (ii) p24-ultrasensitive SIMOA in the supernatant as a proxy for viral particle release (Fig. 6B). While low levels of tat/rev RNA copies were observed in the negative control conditions (medians: 6.5 and 5.4 copies/µg in the non-stimulated and HA-treated condition, respectively), T66 mRNA induced higher levels of latency reversal when compared to PMA (medians: 188 and 76 copies/µg in T66 and PMA-treated cells, respectively). Moreover, while low or undetectable levels of p24 were detected in the negative controls, T66 mRNA induced detectable concentrations of p24 in the supernatant in 6/8 participants. Levels of p24 in the supernatant were slightly lower in the T66 mRNA condition compared to the PMA-treated cells (medians: 185 and 369 fg/mL in the T66 mRNA versus PMA, respectively). In conclusion, the T66 mRNA delivery by the LNP-2 formulation is highly efficient at inducing latency reversal in CD4+ T-cells from cART-treated individuals.

**Figure 6:**
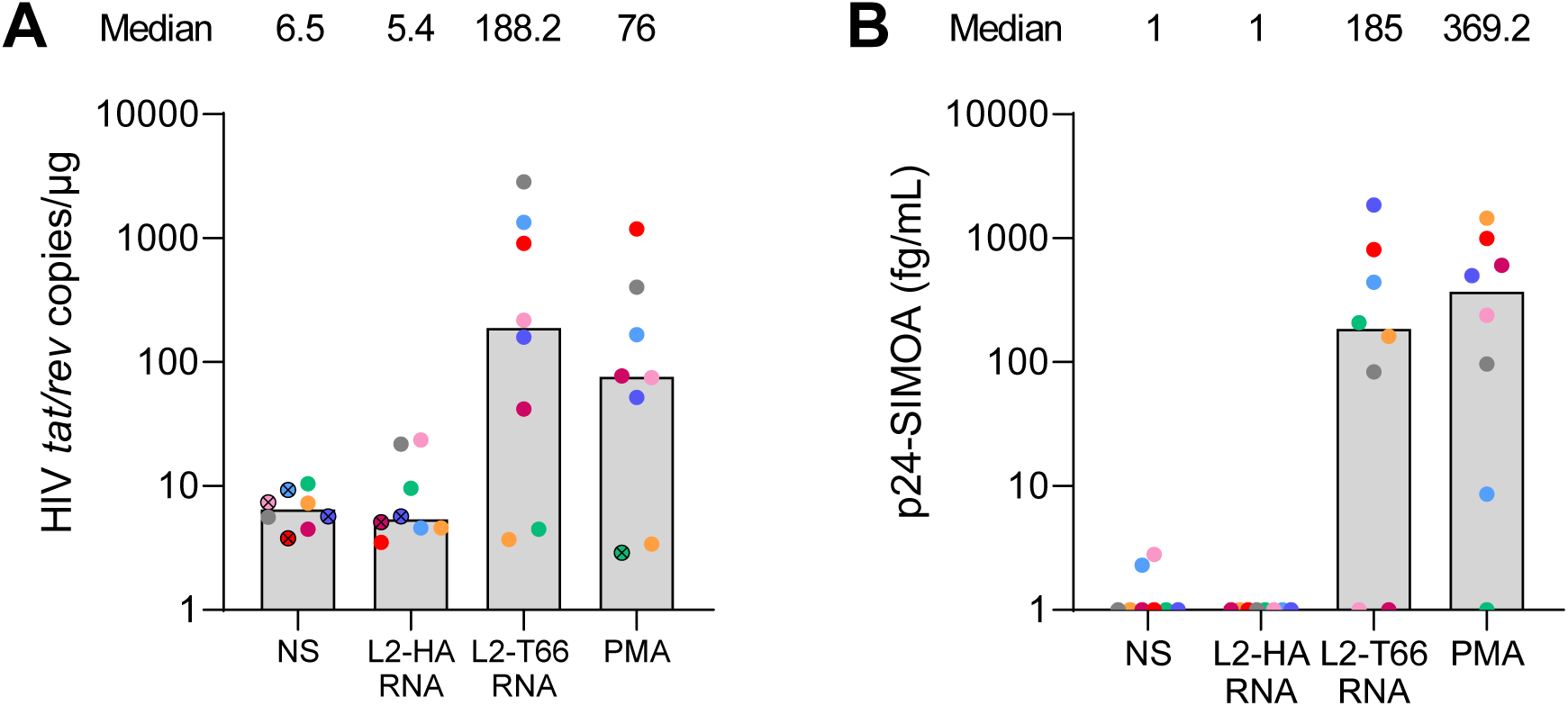
Reactivation of latent infected CD4+ T-cells from HIV infected patients. (A) CD4+ T-cells from 8 different cART treated individuals were treated for 48h with PBS (NS), HA-LNP2, T66 mRNA-LNP2 (Tat LNP2) or PMA as positive control. RNA was extracted from the cells and RT-ddPCR was performed to quantify tat/rev transcripts. Results are expressed as number of *tat/rev* copies per µg of RNA. Undetectable measures are represented by the following symbol ⊗, and limits of detection are plotted. (B) Viral release was evaluated by measuring p24 secretion in supernatant by SIMOA technology.

## Discussion

While cART has greatly improved the quality of life of people living with HIV (Palella et al., 1998), it does not lead to cure (elimination of the virus from its host). The virus persists in a latent, and largely transcriptionally silent state in a small pool of memory CD4+ T-cells (Chun et al., 1998, Finzi et al., 1999) invisible to the immune system. cART treatment interruption studies result in rapid viral rebound in the vast majority of long-term treated HIV-infected individuals(Davey et al., 1999). The latently infected cellular reservoir represents the major roadblock to viral cure. In this paper, we identified a novel agent T66, which is highly efficient at inducing latency reversal, both in cell lines and in primary CD4+ T-cells isolated from cART treated individuals.

HIV-1 Tat protein is probably the strongest transcriptional transactivator known to date (Chun et al., 2015, Ruelas and Greene, 2013). Tat is encoded by two exons: the first exon (72 aa) comprises all essential functional regions and is highly conserved, while the second exon (86-101 aa) codes for a structurally non-defined variable C-terminal region with unknown functionality. The transcriptional effect of Tat is initiated by the binding to TAR, a stem-loop structure in the nascent transcript resulting in the recruitment of the p-TEFb complex to the HIV-1 LTR promoter. This interaction leads to the phosphorylation of RNA Polymerase II (RNAPII) and other associated factors to engage in the production of full-length HIV RNA transcripts from a paused transcription complex (Siliciano and Siliciano, 2013). More recently, it was discovered that p-TEFb also promotes binding of additional elongation factors such as AFF4, ENL, and AF9 to form an assembly called ‘super elongation complex’ explaining the incredible transactivation power of Tat (Chou et al., 2013, He et al., 2010, Sobhian et al., 2010). In the absence of Tat, RNA polymerase II initiates transcription from the LTR promoter by typically stalling downstream of the TAR, producing short, abolished transcripts (Archin et al., 2014, Dahabieh et al., 2015). The ability of Tat to strongly activate viral transcription has long been known, but few attempts have been made to harness the protein as an LRA. The general concern to explore Tat as LRA likely stems from the reported off-target activity and cellular toxicity exerted by full length Tat (Derdeyn et al., 2000, New et al., 1997, Rasmussen et al., 2014, Viscidi et al., 1989). Despite these reports TAR independent effects remain controversial. Transgenic mice expressing full length Tat did not develop any phenotype other than reduced glutathione levels (Choi et al., 2000) or atypical skin lesions after two years of age (Vogel et al., 1988). A more recent study confirmed a total lack of side effects in transgenic mice secreting Tat into the serum (Choi et al.). However, to reduce the risk of potential non-specific receptor engagement by extracellular Tat, we focused on the identification of the smallest variant retaining potent Tat-72 transactivation activity by deletion of the Tat C-terminus which contains the non-structured region of the protein.

The reactivation capacity of the T66 protein was assessed in different cell lines as well as in primary CD4+ T-cells from HIV-infected individuals. The capacity of T66 to reverse HIV latency was shown by three independent laboratories with three different read-outs (cell-associated RNA, p24-SIMOA, and QVOA, (Fig. 3). The cell-associated RNA assay probes for proviral transcriptional activation, the ultrasensitive SIMOA assay confirms that a structural virion protein has been translated and released, and the QVOA assay of resting CD4 T-cells from individuals living with HIV on cART treatment rigorously explores whether replication competent HIV has been produced *ex vivo* by a persistently infected cell.

The T66 protein induced consistently higher levels of cell-associated HIV RNA than the positive control PMA in all tested donors, while it resulted in similar p24 supernatant levels when compared to CD3/CD28 T-cell co-stimulation. However, in the QVOA assay virus was recovered with somewhat greater frequency following mitogen stimulation than T66 treatment. Given the significant heterogeneity in HIV inducibility across different donors generally observed in viral outgrowth assays after LRA exposure, further studies with more donors will be needed to fully assess the *in vitro* latency reversal activity of T66. This is the first time to identify a molecule tested in different HIV latency assays that demonstrates robust induction of HIV expression and latency reversal similar to PKC modulator PMA and importantly in the absence of any CD4 T-cell activation (Spina et al., 2013). The promising activity of T66 merits further testing in combination with other LRAs or immunotherapies *ex vivo* as well as *in vivo* models.

A major barrier to considering Tat as LRA is based on the reported off-target activities and toxicities using recombinant full length Tat86 protein produced in bacteria. Two strategies were applied to address this liability: chemical protein synthesis was chosen over recombinant protein production to avoid any biological contamination with toll like receptor (TLR) ligands that could lead to Pattern Recognition Receptor (PRR) activation of signaling pathways unrelated to Tat-TAR mediated transactivation. Secondly, a large part of the C-terminus was deleted to remove any potential binding sites of Tat to cellular receptors which may be present in this part of the protein such as the RGD motif (residues 78-80) that interferes with some integrin receptors (Chiodelli et al., 2012, Urbinati et al., 2005).

A major clinical challenge will be to deliver sufficient T66 protein to HIV-infected CD4+ T-cells *in vivo*. Administration of a substantial amount of protein may be required due to inefficient cellular protein transduction, significant endosomal trapping, high plasma protein binding, and potential neutralization by preexisting anti-tat antibodies. One strategy to address these possible protein-associated limitations would be to deliver T66 mRNA packaged into nanoparticles to CD4+ T-cells. A proof-of-concept study *in vitro* was initiated to evaluate whether T66 mRNA would induce similar HIV reactivation as observed with the T66 protein. Most notably, T66 mRNA transfected with jetMessenger into a MT4 HIV-GFP reporter cell line exceeded the activation of T66 protein by more than 4 log_10_. Maximal reporter expression was achieved with 0.5nM T66 mRNA while it required 5 µM of protein to reach similar activation. Finally, the ability of T66 mRNA to induce latency reversal in a difficult to reactivate Jurkat clone 50 cell line was assessed: 6 nM of T66 mRNA transfected with j jetMessenger showed equipotency with PMA. The recognition of superior T66 mRNA activity led to test additional engineered LNPs for efficient mRNA delivery into primary lymphocytes. LNP-2 was selected as the lead LNP that significantly exceeded the level of reactivation observed with the jetMessenger formulation. The LNP-2 formulation was therefore further explored in CD4+ T-cells from HIV-infected individuals. T66 mRNA delivered by LNP-2 induced higher levels of *tat/rev* mRNA than PMA stimulation, and slightly lower levels of p24 release in the supernatant compared to PMA.

In conclusion, our work suggests that T66 mRNA shows similar potency to the gold standard activation compound PMA to induce latency reversal and viral particle release. In contrast to PMA, T66 exposure to primary CD4+ T cells led to neither T cell activation nor showed any significant perturbation of the human transcriptome. Therefore, T66 protein or mRNA could be an important LRA to be considered for HIV cure combination regimens. Clinical translation of our approach will require validation of the latency reversal capacity and the absence of toxicity in a relevant preclinical model.

## Materials and methods

### Participants and blood collection

In this study, patients living with HIV (PLWH) on stable suppressive cART were recruited from four different institutes: Institute of Tropical Medicine (Antwerp, Belgium; n=14; Cohort A), Hospital Pitié Salpetière (Paris, France; n=10; cohort B), University of North Carolina HIV Cure Center (United States; n=8, cohort C), and HIV Cure Research Center (Ghent, Belgium; n=8, cohort D). HIV-infected individuals included in this study were all on cART for at least 1.5 years and had a viral load <20 copies/mL. Patient characteristics are summarized in Table 1. Peripheral blood mononuclear cells (PBMCs) from healthy donors were obtained from the Red Cross (Mechelen, Belgium).

**Table 1:** patient characteristics

| Laboratory | Cohort A | Cohort B | Cohort C | Cohort D |
| --- | --- | --- | --- | --- |
| Age in years, median, (IQR) | 52(34-69) | 50(34-58) | 46(31-58) | 52 (48-62) |
| CD4 in cells/mm<sup>3</sup>median (IQR) | 821(361-1113) | 628(462-689) | 636(525-797) | 823(335-126) |
| CD8 in cells/mm<sup>3</sup>median (IQR) | 1113(369-1689) | 876(643-10007) | 513(414-632) | 963(567-1800) |
| CD4/CD8 ration median (IQR) | 0.67(0.31-2) | 0.7(0.6-1) |  | 1(0.5-1.3) |
| CD4 nadir in cells/mm<sup>3</sup>median (IQR) | 197(83-285) | 241(209-300) | 388(119-532) | 291(98-488) |
| Years since first diagnosis median (IQR) | 17.3(3.6-24) | 19(25-10) | 4.58(2-17.9) | 16(8.4-30.1) |
| Years on cART median (IQR) | 15.7(3.4-17) | 7.2(5.1-11.0) | 4.40(1.9-15.6) | 12(1.2-24.3) |
| Viral load log10 (copies/ml) | <20 | <20 | <20 | <20 |
| Years suppressed median (IQR) | 12(1.2-24.3) | 7.2 (5.1-11) | 3.56(1.8-7.1) | 15.7(3.4-17) |

### Ethics statements

All participants were adults and signed informed consent forms approved by the respective institutional Ethics Committees.

### Generation of reporter cell lines to measure latency reversal activity

HEK293T cells (ATCC, Manassas, USA) were transfected with pNL-EGFP/pVSV using polyethylenimine (PEI; Polysciences, Germany) (PEI/DNA 6/1 ratio; 500 µL/well). 48h post transfection virus was collected and concentrated with PEG Virus Precipitation Kit (BioVision Inc, USA) according to the manufacturer’s instructions. The Jurkat Clone E6.1 (ATCC, USA) cell line was then infected with PEG concentrated virus (1/10 ratio) for 24 h. After removing the virus, the cells were cultured for 1 week. The GFP (-) cell population potentially harboring latent virus was sorted and stimulated with 10 ng/mL Phorbol 12-Myristate 13-Acetate (PMA, Sigma Aldrich) followed by sorting the GFP (+) population. After 1 week the GFP negative cells were sorted again and restimulated with 10 ng/mL PMA. This process was repeated 4 times. Finally GFP(+) were single cell sorted and cultured for 3 weeks. After this process 2 clones were selected with low GFP background, Jurkat E6.1 clone 32 and Jurkat E6.1 clone 50), which contain latent HIV with a GFP reporter.

To generate MT-4-LTR-EGFP cells, MT-4 cells (gift from(Nakashima et al., 1990) were transfected with a plasmid containing LTR-driven EGFP expression plasmid (Clontech Laboratories Inc, USA) and pFX2 (Invitrogen Corporation, USA) in Opti-Mem Reduced Serum Media (Thermo Fisher Scientific Europe) according to the manufacturer’s instructions. Cells were selected based on geneticin (Thermo Fisher Scientific, Europe) resistance. To obtain a stable cell line, cells were sorted based on high viability and low GFP background.

### Generation of plasmids

The full-length Tat protein (86-101 aa) is translated from two different exons where exon-1 (1-72 aa) contains all domains essential for trans-activation. The following Tat coding sequences (based on the molecular HIV clone NL4.3) were introduced into the DNA expression vector pVax-1 (Life Technologies): tat86 (86 referring to length in amino acids) *tat*72, *tat*70, *tat*68, *tat*66, *tat*65, *tat*64, *tat*60, *tat*57. The HIV-1NL4.3 LTR was cloned into pG5luc (Promega) upstream of the firefly luciferase gene, yielding the plasmid pLTR-FLuc. The transfection normalization plasmid pEF-RLuc was generated by swapping the CMV promoter with the EF1α (Elongation Factor 1 α) promoter in pcDNA3.1 followed by the downstream introduction of the Renilla luciferase coding sequence in the multiple cloning site.

### Dual Luciferase assay for Tat activation assessment

HEK293T-cells were co-transfected with the different *tat* expression plasmids, the pLTR-FLuc, and the pEF-RLuc plasmids. The Dual Glo-reporter assay (Promega) was used to measure LTR activation via the Firefly luciferase readout along with signal normalization by Renilla luciferase. LTR activity was expressed as percentage of wild type exon-1 *tat*72 activity.

### Production of the T66 protein

The Tat66 (T66) protein was synthesized as trifluoroacetate salt by Bachem. The provided lyophilized powder was resuspended in sterile water containing 1mM DTT to a final concentration of 1 mg protein/mL. The pH was adjusted with NaOH to pH 7.4.

### HIV reactivation in cell lines

MT-4-LTR-GFP, Jurkat E6.1 clone 32 and clone 50 were incubated overnight with a dose range of the T66 protein or T66 mRNA. Medium was used as a negative control, while PMA (10 ng/mL), CD3/CD28 Dynabeads (1 bead per cell, Thermofisher, Europe) or Phytohemagglutinin (10 µg/mL; PHA, Sigma Aldrich) were used as positive controls. LTR activation was measured by flow cytometry (BD Fortessa, BD biosciences, Erembodegem, Belgium) based on GFP expression.

### HIV reporter assay on primary CD4+ T-cells

PBMCs were isolated from HIV-negative blood donors by Ficoll gradient centrifugation. CD4+ T-cells were purified by positive selection using Miltenyi magnetic beads (Miltenyi Biotec, Gladbach, Germany). These cells were stimulated with anti-CD3/CD28 magnetic Dynabeads (1 bead per cell, Thermofisher, Europe) for 2 days. Following the removal of the beads, cells were co-infected with a single cycle HIV dual reporter virus expressing GFP and mouse CD48 and a lentiviral vector coding for pro-survival proteins MCL1 and Bcl2 co-expressing cell surface mouse CD24. Infected cells were cultivated in the presence of IL-2 (10 IU/mL, Roche, Basel, Switzerland). Three days after coinfection cells were sorted based on expression of mouse CD48 and mCD24 (BD biosciences, Belgium). These cells were further cultivated for another 7 days on IL-2 containing medium and rested for 3 extra days in assay medium (RPMI, 10% FCS). To evaluate HIV reactivation cells were plated at a concentration of 2 million per mL and incubated in the presence of T66 protein (4 µM), negative control (PBS) or positive control PMA (5 ng/mL) for 24h. Reactivation capacity was evaluated by measuring the percentage of GFP+ cells using flow cytometry (Fortessa, BD Biosciences, Belgium).

### Ex vivo reactivation of HIV measured by RT-qPCR

CD4+ T-cells were isolated from 200 mL whole blood donated by cART treated individuals, using CD4 microbeads (Miltenyi Biotec, Germany) according to manufacturer’s protocol. Resting CD4+ T cells were isolated from leukapheresis products as previously described(Archin et al., 2017).Ten to twenty replicates of cell pools plated at 1 million total or resting CD4+ T--cells per well were incubated overnight in media (RPMI, 10%FCS) with the T66 protein (4 µM), negative control (PBS) or positive control (PMA, 5 ng/mL or highly purified PHA 1.5µg/ml Thermofisher, Europe). Total RNA was isolated using the MagMax 96 total RNA isolation kit (Ambion, Europe) following the manufacturer’s protocol. Reverse transcription was performed using the Superscript III First Strand synthesis kit (Invitrogen, Europe) according to the manufacturer’s protocol. Quantitative real time PCR (qPCR) was conducted on the cDNA using gag-specific primers (GGACCAAAGGAACCCTTTAGAGA; GGACCAACAAGGTTTCTGTCATC) in the presence of nucleic acid dye SYBR Green (Invitrogen, Europe). Standard curves were generated using cDNA synthesized from *in vitro* transcribed RNA. The detection limit of the qPCR was determined to be at 10 copies/reaction. Relative HIV-1 gag RNA copies were calculated related to the standard curve.

### Ex vivo reactivation of HIV measured by p24 detection

PBMCs were isolated from whole blood of cART suppressed individuals by Ficoll gradient centrifugation. CD4+ T-cells were purified from PBMCs by negative selection (EasySep ™ Human CD4+ T-cell Isolation Kit, Stem Cell, Cambridge, UK). CD4+ T-cells were plated in 200ul of medium (RPMI, 10% human serum) containing IL-2 (Roche, 10 IU/mL) and IL-7 (Peprotech, 1 µg/mL) in triplicate (200,000 cells/well). The T66 protein was added to a final concentration of 4 µM. Anti-CD3/CD28 antibodies were used as a positive control, while PBS was used as a negative control. Cells were incubated at 37 °C for 10 days and medium was changed every 3-4 days (180 µL of supernatant removed and refreshed with new medium supplemented with IL-2 and IL-7). Supernatant was frozen at −80 °C for subsequent detection of p24 by SIMOA (Quanterix, USA). In another set of experiments, CD8 T-cells were depleted from the PBMC by positive selection (EasySep^TM^ CD8 T cell isolation kit, Stem Cell, Cambridge, UK). CD8-depleted PBMCs were cultured in medium (RPMI, 10% human serum) and treated as described above for the CD4+ T-cells. Frozen supernatant from above cultures were thawed and p24 production was evaluated by SIMOA following the manufacturer’s instructions.

### Ex vivo reactivation of HIV measured by qVOA

The qVOA assay was performed as previously described (Archin et al., 2017, Crooks et al., 2015). Briefly, 34-50 million resting CD4+ T-cells were plated in replicate (n=8) at limiting dilutions (2.5, 0.5, 0.1 and 0.025 million cells per well), activated with PHA (Remel, Thermofisher, US) in the presence of a fivefold excess of allogeneic irradiated PBMCs from an HIV seronegative donor and 60 IU/ml IL-2 or T66 protein (4µM) for 24h. Afterwards cells were co-cultivated with CD8-depleted PHA blasts. Culture supernatants were harvested on day 15 and assayed for virus production by p24 antigen capture ELISA (ABL, Rockville, MD, United stated). A maximum likelihood method was used to estimate the frequency of replication-competent proviruses, reported as infectious units per million of CD4+ T-cells (Myers et al., 1994, Trumble et al., 2017).

### Transcriptome analysis by micro-array

Bulk CD4+ T-cells isolated from cART treated individuals were treated with T66 protein (4µM), PHA (10µg/mL) or PBS. Twenty-four hours later 100,000 cells were transferred to a tube containing 100 µl of RLT buffer (Qiagen, Hilden, Germany). Cells were stored at –80 °C until further processing. RNA extraction was prepared with the RNeasy plus mini kit (Qiagen, Germany). Amplification and labelling of total RNA were performed using the GeneChip® PICO Reagent Kit following the manufacturer’s protocol (ThermoFisher, Europe). Biotin-labeled target samples were hybridized to the Clariom™ GO Screen containing probes for over 20,000 genes. Target hybridization was processed on the GeneTitan® ™ Multi-Channel Instrument (Applied Biosystems™, Waltham, USA) according to manufacturer’s instructions provided for Expression Array Plates. Images were analyzed using the GeneChip® Command Console Software (GCC) (ThermoFisher, Europe). The microarray dataset was background corrected and normalized with robust multiarray analysis [38]] and summarized with the ClariomSHumanHT_Hs_ENTREZG v21.0.0 chip definition files [39]. Quality assessment was performed with array QualityMetrics (Kauffmann et al., 2009). Using the expression values for all 22,593 probe sets, we evaluated the main source of variability contained in the dataset using principal component analysis (PCA). We performed differential gene expression by comparing different treatments) using linear regression models for microarray data analysis (Limma) (Smyth, 2004).

### Evaluation of global T-cell activation by flow cytometry

Following treatment with the T66 protein (4 µM), PMA (8 nM) or the negative control (PBS) for 24 h, CD4 T-cells were stained with anti-CD25 PerCp-cy5.5 and anti-CD69 APC (BD biosciences, Belgium) for 15min at 4 °C. Cells were analyzed on Fortessa (BD Biosciences, Belgium).

### Generation of T66 mRNA

A DNA template for the *in vitro* transcription of mRNA was generated by introducing the following elements in a standard pDNA3.1 vector (Thermofisher, Europe): a T7 promoter, followed by 5’-and 3’-UTRs from either the human α-globin (HAG) or the frog (Xenopus) α-globin (XBG), terminated by a 120 nt poly-A tail. Downstream of the Poly-A tail is a unique restriction enzyme site for linearization. In reactions using the capping reagent AG CleanCap (Trilink, San Diego, USA) the native T7 promoter sequence was changed to TAATACGACTCACTATAAG. An HIV-1 T66 codon-optimized gene sequence was introduced via restriction enzyme-based cloning into the DNA plasmid between the 5’-UTR and 3’-UTR. Sequence-verified DNA constructs were scaled-up, linearized and column-purified with the PureLink Gel Extraction Kit (Thermofisher, Europe). *In vitro* transcription was performed with either HiScribe T7 ARCA mRNA kit or the HiScribe *In Vitro* Transcription Kit (NEB) according to the manufacturer’s recommendation. 500 ng linearized plasmid DNA (pDNA) was added per 20 µL transcription reaction. The ribonucleotide uridine was in some cases replaced with N1-methypseudouridine. The capping reagent AG Cleancap (Trilink, USA) was added at 5mM concentration to the transcription reaction along with 0.1 U of inorganic pyrophosphatase YIPP (NEB). *In vitro* transcription was typically performed for 3 h at 37 °C. The *in vitro* transcript was purified by column purification using RNA miniprep/maxiprep kits (Qiagen, Germany) according to the provided protocol. The RNA was quantified with the Nanodrop spectrophotometer, and its integrity verified by agarose gel analysis.

### Identification of nanoparticle formulations for efficient T66 mRNA delivery

A panel of commercial transfection reagents were tested in a cellular HIV-GFP reporter assay to identify the optimal delivery system of T66 mRNA into Jurkat T lymphocyte cells: JetPEI (Polyplus, Illkrich-Grafenstaden, France), JetMessenger (Polyplus, France), Viromer (Lipocalyx, Halle, Germany), TransIT mRNA (Mirus, Marietta, USA), RNAiMax (Thermofisher, Europe), and Messenger Max (Thermofisher, Europe). The transfections were performed with 1×10^5^ Jurkat cells plated in 100 µL medium in 96-well plates using 100 ng/ml T66 mRNA formulated with the different reagents according to the supplier’s instructions.

Lipid nanoparticles (LNPs) were prepared by mixing a lipid/ethanol solution with an aqueous RNA solution. Specifically, lipid excipients (ATX—Arcturus Therapeutics proprietaryionizable lipid, phospholipid, cholesterol, and polyethylene glycol derivatized lipid) were dissolved in ethanol. An aqueous solution of the RNA was prepared in citrate buffer (pH 3.5). The lipid mixture was then combined with the RNA solution using a NanoAssemblr microfluidic system (Precision NanoSystems). Nanoparticles thus formed were dialyzed against HEPES buffer (pH 8.0) using dialysis tubing (Repligen, Waltham, USA) at room temperature. The concentration of the formulation was adjusted to the final target RNA concentration using Ultra centrifuge concentrator tubes (Millipore Sigma, Burlington, USA) and was sterile filtered. Post filtration, bulk formulation was aseptically transferred into sterile vials and frozen at −70 ± 10 °C. Analytical characterization included measurement of average particle diameter and degree of size heterogeneity of LNPs (ZEN3600, Malvern Instruments), RNA content, and encapsulation efficiency by a fluorometric assay using Ribogreen RNA reagent (Thermo Fisher Scientific).

### Ex vivo reactivation of HIV in primary CD4 T-cells by T66 mRNA

Two million CD4+ T-cells, isolated from PBMC of cART treated individuals, were stimulated for 48h with T66 mRNA packaged in LNP-2 (250 ng/mL, Arcturus) in presence of antiretroviral drugs (200 nM raltegravir, 200 nM lamivudine). A 24h-stimulation with PMA (5 ng/mL) was included as positive control, while non-stimulated cells and cells incubated with HA LNP-2 (lipid nanoparticle formulation containing the hemagglutinin peptide from influenza) were used as negative controls. Following stimulation, cells were centrifuged (2,500 rpm, 10 min) and supernatant was collected for assessment of p24 release by p24-SIMOA (the protocol is described above). Cells were centrifuged at high speed (14,000 rpm, 10 min) and dry pellets were stored at –80 °C until further processing. RNA extraction using the innuPREP RNA Mini Kit (Westburg, #AJ 845-KS-2040250) yielded a median concentration of 48.6 ng/µL (elution in a final volume of 30 µL). The reverse transcription (RT) and digital droplet PCR (ddPCR) steps were performed as described previously (Yukl et al., 2018). In brief, the RT step was done in a final volume of 25 µL and the mix was composed of: 2.5 µL 10X Superscript III buffer (Invitrogen #10308632), 2.5 µL MgCl2 50 mM, 1.25 µL random hexamers 50 ng/µL (Invitrogen #10308632), 1.25 µL dT15 50 µM, 1.25 µL dNTPs 10 mM, 0.625 µL RNAse OUT 40 U/µL (Invitrogen #10308632), 1.25 µL SuperScript III RT 200 U/µL (Invitrogen #10308632) and 14.4 µL of extracted RNA. Thermocycling conditions were as follows: 10 min at 25 °C; 50 min at 50 °C; 5 min at 85 °C. ddPCR on *tat/rev* transcripts was performed using the QX100 Droplet Digital qPCR System (Biorad). The 20 µL PCR mix was composed of: 10 μL ddPCR Probe Supermix (no dUTP), 1.8 μL *tat/rev* primers 900nM (F: 5’-CTTAGGCATCTCCTATGGCAGGAA-3’, R: 5’-GGATCTGTCTCTGTCTCTCTCTCCACC-3’), 0.5 μL probe 250 nM (5’-ACCCGACAGGCC-3’), 0.9 μL H2O and 5 μL undiluted RT product. Droplets were amplified (Thermal Cycler T100, Bio-Rad) using the following cycling conditions: 10 min at 95 °C, 45 cycles (30 sec at 95 °C, and 60 sec at 59 °C), 10 min at 98 °C. Each sample was measured in triplicate.

### Statistical analysis and software used

The Wilcoxon rank sum test was used to calculate the statistical significance of the relative gag RNA copy number between different conditions. For micro-array analysis genes identified were considered statistically significant based on Benjamini-Hochberg adjusted p-value <0.05.

We used R software, version 3.6.1 (2019-07-05), platform x86_64-pc-linux-gnu (64-bit), running under: Red Hat Enterprise Linux Server 7.9 (Maipo) and additionally utilized R packages, including Complex Heatmap (Gu et al., 2016) for data visualization. GraphPad Prism software (v9.0) was used to visualize data and perform statistical analysis.

## Acknowledgments

We thank all participants who donated blood samples, as well as MDs and study nurses who helped with the recruitment and coordination of this study. Part of this research work was supported by VLAIO O&O (HBC.2018.2278). MP was supported by postdoctoral funding from VLAIO O&O (HBC.2018.2278), work at UNC was funded by CARE.  We also would like to thank Jef Hens and Matias Van Den Berck from Charles River Laboratories (Beerse, Belgium) for contributing to this work and Jinho Park from Arcturus Therapeutics for formulating the LNP.

## Author contributions

EVG, DB, MP conceptualized the experiments. EN, NV, KD, CVDE, EB have developed, performed, analyzed the ex vivo reactivation assay, p24 SIMOA, and flow cytometry. CH performed the experiments on T66 protein. MP performed the experiments evaluating T66-LNP in CD4 T-cells from cART treated individuals. IVDW and FE performed and analyzed the micro-array data. NA and DMM were responsible for executing and analyzing additional SIMOA experiments, the QVOA assay, and obtaining human subjects approvals at the University of North Carolina at Chapel Hill. EF obtained ethical approval and was responsible MD for samples recruited at ITM. BA and KK are the responsible MD and requested ethical approval for studies performed at Hospital Pitié Salpetière. LV is the responsible MD who requested ethical approval at Ghent University. EVG, MP and DB wrote the paper. All authors read and edited the paper.

