## supplementary figure 1 for "A truncated HIV Tat demonstrates potent and specific latency reversal activity"

**Supplementary Figure legends**

**Supplementary figure 1: T66 protein reactivates latently infected CD4+ T-cells as measured by cell-associated RNA** CD4+ T-cells from 4 cART treated individuals were exposed 24hours to PBS, T66 protein, or PHA. HIV-1 reactivation following latency reversal was assessed by RT-qPCR and presented here normalized to PHA activity.

**Supplementary Figure 1**

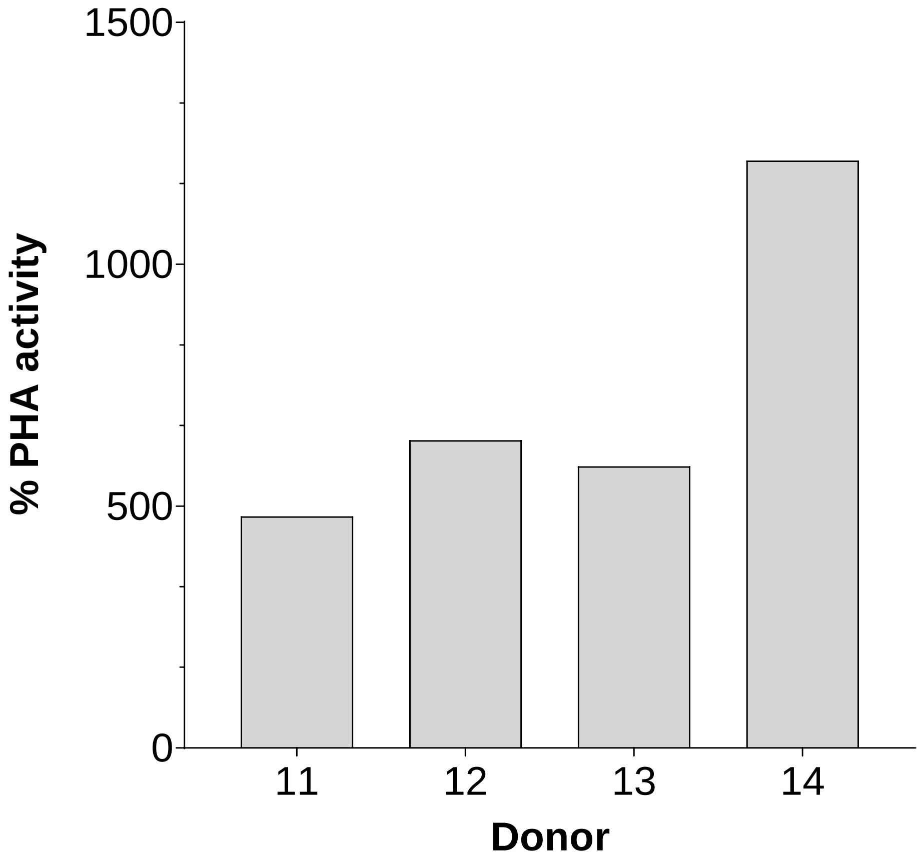
